## Supplementary Figures for "Sensory coding and causal impact of mouse cortex in a visual decision"

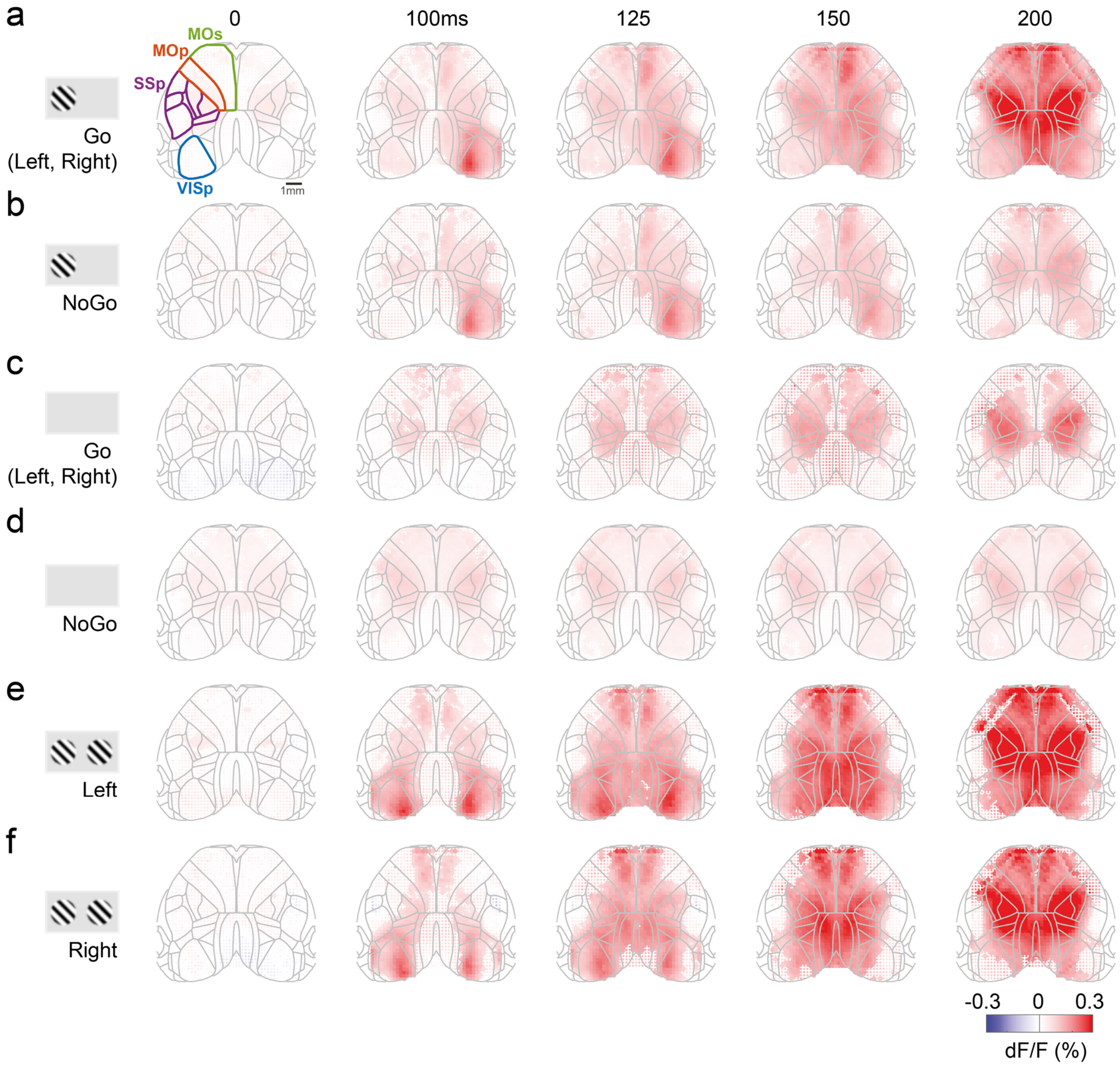


**Figure 2–figure supplement 1: Average cortical fluorescence (dF/F) for different task conditions. (a)** Timecourse of cortical fluorescence, averaged over Go trials (Left or Right choice), with positive contrast present on the left side, and zero contrast on the right. Columns indicate successive time steps after stimulus onset. Each map is overlaid with an outline of cortical regions defined by the Allen CCF, and widefield fluorescence is cropped to the outer edges. Four different cortical regions are highlighted in the left panel (VISp, SSp, MOp, MOs). Maps are averaged over 39 sessions in 9 mice, with dot size indicating significance across sessions ($p$<0.001; nested ANOVA). Even though the animal turns the wheel, the stimulus does not move on the screen in this time period (see Figure 2-figure supplement 2a). **(b)** Average cortical fluorescence for NoGo trials with positive contrast on the left screen. The maps in (a) and (b) are averaged over trials with matched contrast values **(c)** Average cortical fluorescence for Go trials (Left or Right choice) with zero contrast on both sides. **(d)** Average cortical fluorescence for NoGo trials with zero contrast on both sides. **(e)** Average cortical fluorescence for Left choice trials with equal non-zero contrast on both screens. **(f)** Average cortical fluorescence for Right choice trials with equal non-zero contrast on both screens, and mice made a Right choice. The maps in (e) and (f) are averaged over trials with matched contrast values. Panels a,b,e,f are reproductions of Fig. 2b-e, added here for completeness.

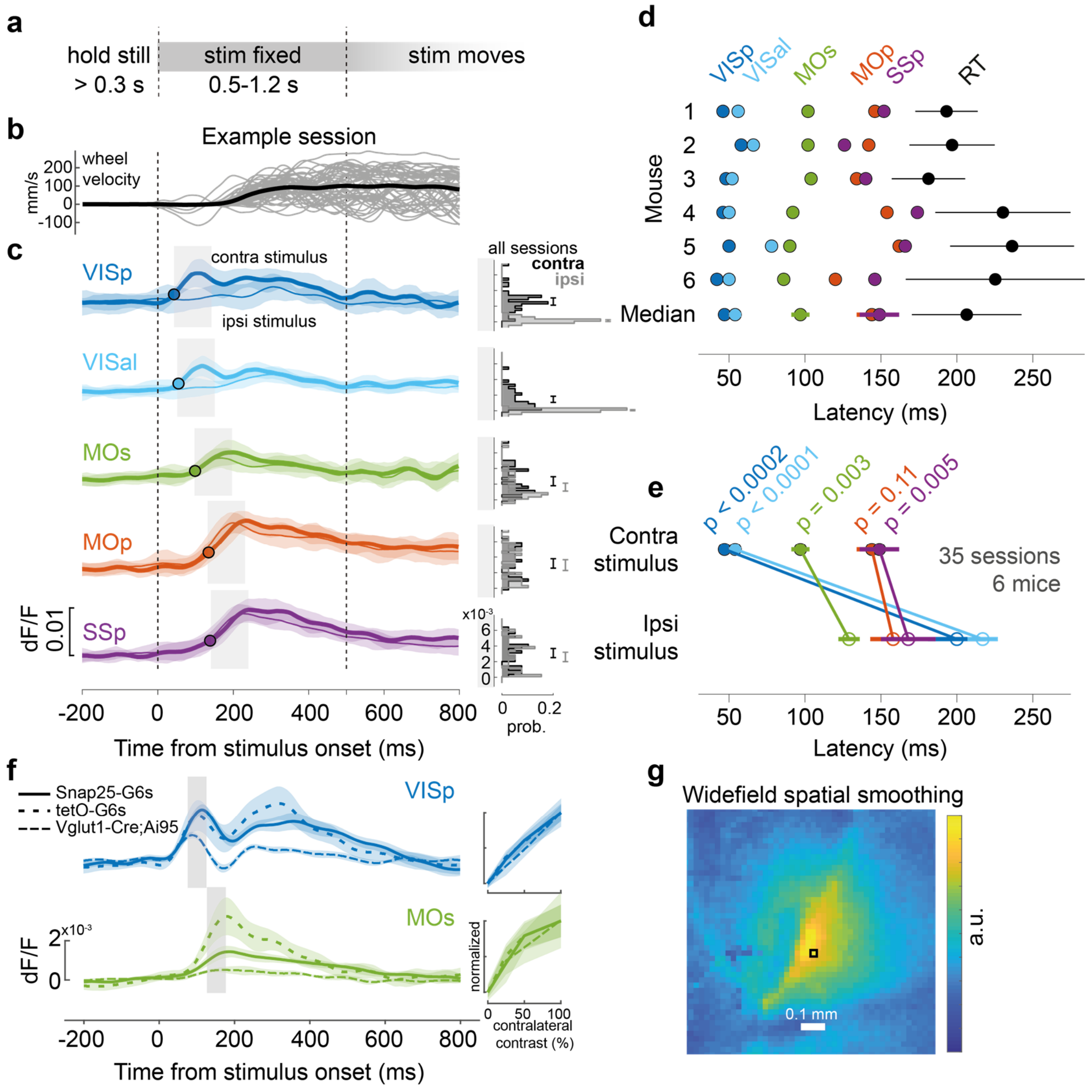


**Figure 2–figure supplement 2: Widefield sequential activation. (a)** Task timeline during widefield calcium imaging. To distinguish activity associated with initial wheel movement from activity driven by visual motion, we introduced an open-loop period 0.5-1.2 s after grating onset, when wheel movements did not move the grating. Trials were excluded post-hoc if choices were made after this period. **(b)** Stimulus-locked wheel velocity for one example session. Thin lines are individual trials for all correct Left or Right choices (velocity sign flipped for Left choices so that correct is always positive), smoothed with a 10 ms Gaussian window. Thick line: mean. **(c)** Stimulus-triggered mean calcium fluorescence at 5 ROIs for one example session. Thick lines: mean response to correct trials with only contralateral stimuli. Thin lines: same for correct trials with only ipsilateral stimuli. Shaded regions: standard deviation of fluorescence across trials. Black circle: response latency, defined as time for mean fluorescence to reach 30% of the peak. Right: distribution of calcium fluorescence for each ROI across all sessions, taken over a 100 ms time window aligned to response latency (shaded regions on left), separated for contralateral (black) and ipsilateral (gray) stimulus trials. Error bar is 95% confidence interval for the mean fluorescence. **(d)** Summary of fluorescence response latencies to contralateral stimuli for 6 mice (see Methods for genotype information; 3 mice were excluded due to having too few trials to reliably measure onset latency). Rows show data from individual mice, with dots showing each ROI's response latency for trials pooled across sessions. Bottom row: response latencies and reaction time (RT) averaged across mice (dots and lines: median ± m.a.d.). **(e)** Response latencies to contralateral and ipsilateral stimuli for each region, averaged across mice (closed and open circles). Significance was determined by a two-tailed paired t-test, $t(5)$=-10.38, -12.00, -5.46, -1.97, -4.72. Colors as in (c). **(f)** Stimulus-locked calcium fluorescence at VISp (blue) and MOs (green) ROIs for three different GCaMP6 mouse genotypes (see Methods). Lines indicate the session-averaged fluorescence for trials with only contralateral stimuli, pooling sessions from Snap25-G6s (solid), tetO-G6s (light dashed), Vglut1-Cre;Ai95 (heavy dashed) mice. Shaded regions indicate the 95% confidence interval. Right inset: fluorescence averaged over time windows depicted by the gray bars on the left, separated for different contralateral contrast values and normalized to the same maximum and minimum across all genotypes. **(g)** Widefield data were

preprocessed by singular value decomposition (SVD), which compresses, smooths, and denoises the calcium signals (Peters et al., 2021). The spatial smoothing that SVD causes depends on the signal statistics and is not spatially homogeneous or isotropic. This pseudocolor image shows the equivalent smoothing kernel resulting from SVD preprocessing, i.e. the SVD-smoothed imaging that would result from a raw image with a single pixel over Right VISp (black square), and zero everywhere else. The kernel has ~0.1-0.2 mm diameter. White: scale bar.

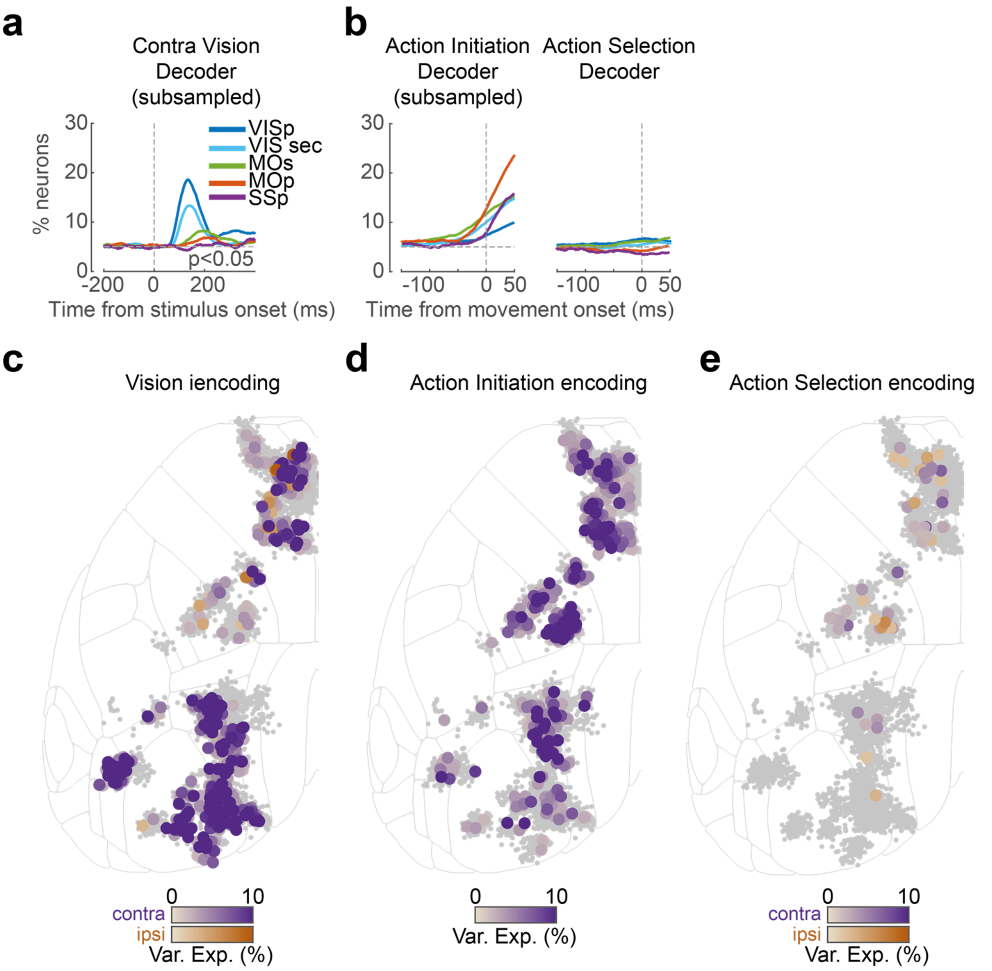


**Figure 2–figure supplement 3: Neuropixels decoding analysis and encoding (kernel regression) analysis. (a,b)** Same binary decoding results as in Fig. 2m-p but using a subsampled number of trials to match with the choice decoder. VIS sec represents the signal from multiple secondary visual areas. **(c)** Enlarged version of Fig. 2j, showing single neurons with significant cross-validated variance explained using a contralateral (purple) or ipsilateral (orange) stimulus kernel. **(d)** Enlarged version of Fig. 2k, showing single neurons with significant movement kernels **(e)** Enlarged version of Fig. 2l, showing single neurons with significant kernels for contralateral (purple) or ipsilateral (orange) choice.

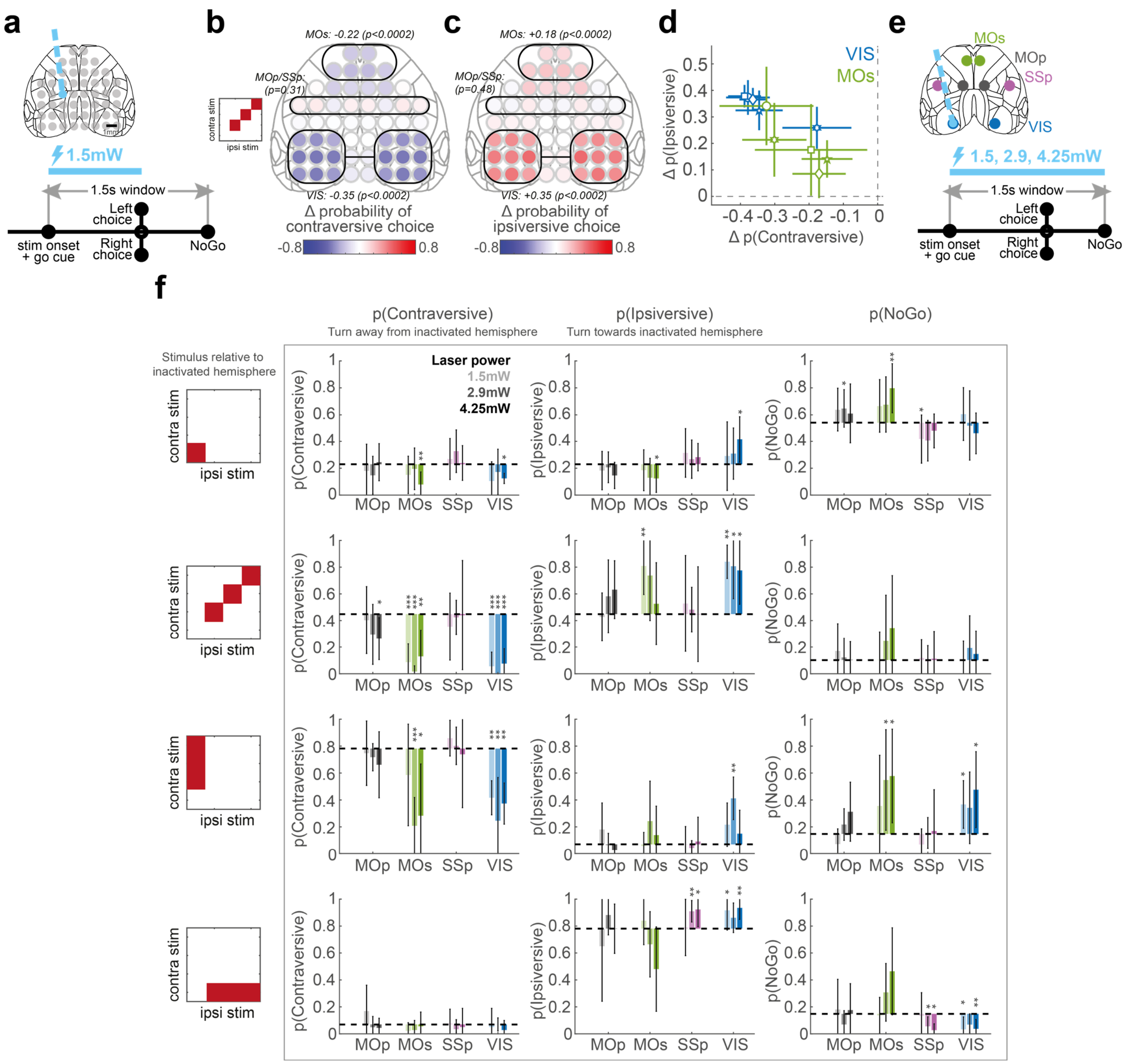


**Figure 3–figure supplement 1: 52-coordinate and mixed-power optogenetic inactivation. (a)** Schematic of the 52-coordinate inactivation experiment (91 sessions in 5 mice). On ~75% of trials, a laser (473 nm, 40 Hz, 1.5 mW) was switched on during stimulus presentation and ceased when a choice (or NoGo) was registered. The location of the laser varied randomly trial-to-trial over 52 cortical sites. **(b)** Summary map of the effect of laser inactivation on contraversive choices, on trials with equal non-zero contrast on each side. Colors indicate the change in the probability of making a contraversive choice (Left choices for inactivation of right hemisphere locations, and Right choices for left hemisphere locations), averaged across 91 sessions in 5 mice. Data are plotted symmetrically across the hemispheres. Black lines mark regions for which statistical significance was assessed (black italic text) by pooling these regions and shuffling the identities of laser and non-laser trials within each session. **(c)** Summary map of inactivation effect on ipsiversive choices, plotted as in (b). **(d)** Mouse-to-mouse variability in inactivation results. Each marker represents the mean change in contraversive and ipsiversive choices for a single mouse on inactivation of any site in VIS (blue) or MOs (green). Each mouse represented by a distinct glyph shape. Error bars represent 95% confidence interval. Dashed gray lines: 0 change in choice proportions. **(e)** Schematic of mixed-power fixed-duration inactivation (34 sessions in 6 mice), focused on VIS, MOs, MOp and SSp. Inactivation was performed at several laser powers (1.5, 2.9, 4.25 mW), and the inactivation duration was fixed at 1.5 s, starting at visual stimulus onset. **(f)** The behavioral effect of inactivation on Contraversive, Ipsiversive and NoGo responses (columns) for zero contrast, bilateral, and unilateral stimulus conditions (rows), separated for the different laser powers (shading). Movement direction and stimulus side is expressed with respect to the inactivated hemisphere. Bars indicate the mouse-averaged choice proportions, and error bars indicate 95% confidence intervals on the mean estimate. Dashed gray lines indicate

the average choice proportions with no inactivation. Statistical significance across mice was determined using a t-test. Significance results are not corrected for multiple comparisons. *** $p<0.001$, ** $p<0.01$, * $p<0.05$.

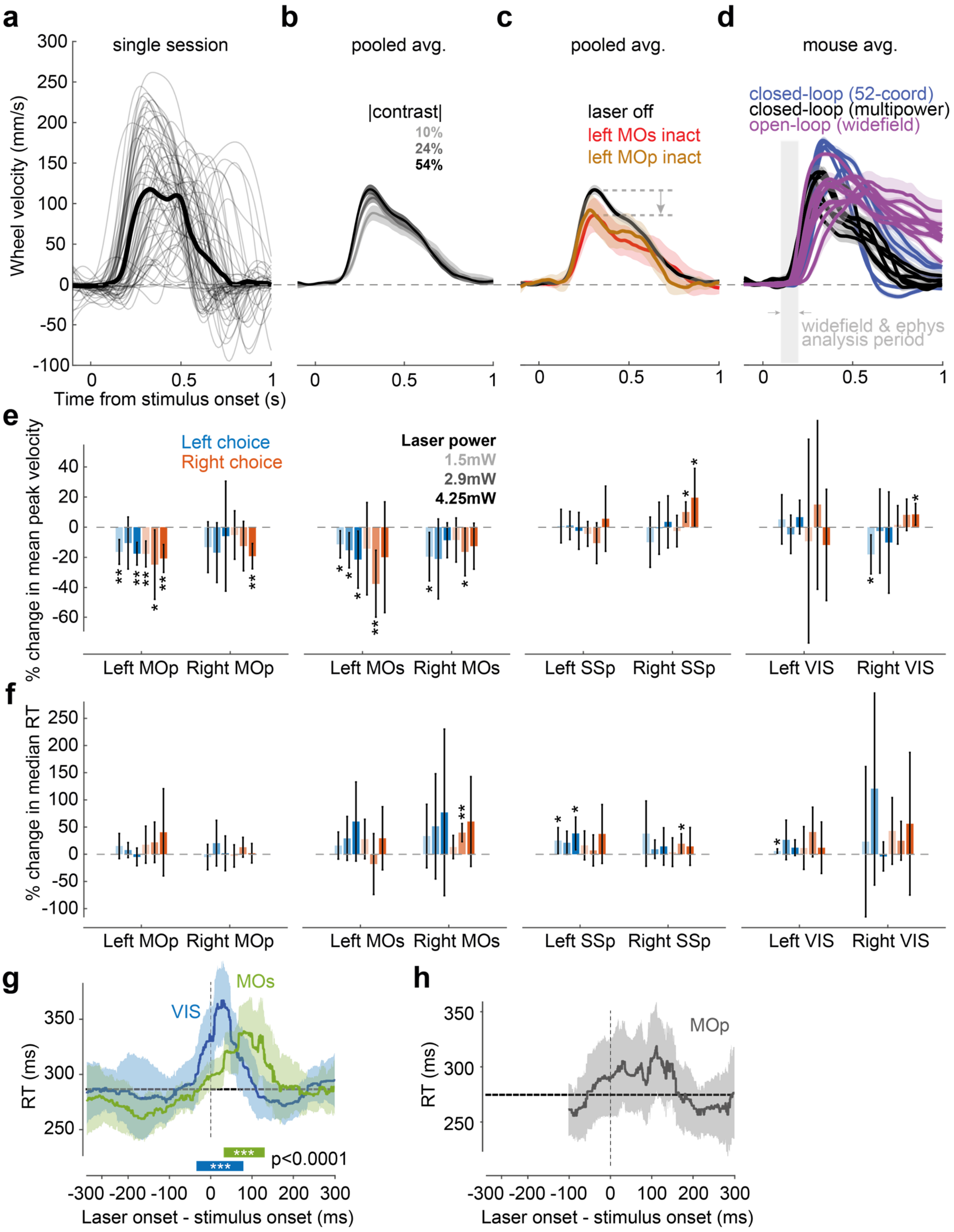


**Figure 3–figure supplement 2: Effect of visual contrast and optogenetic inactivation on wheel movements. (a)** Wheel velocity for a single session, for all correct choice trials with a single stimulus on one side (velocity sign flipped for Left choices so that correct is always positive). Thin lines: single trials smoothed with a 10 ms Gaussian window. Thick line: mean across trials. **(b)** Mean wheel velocity pooled over all correct non-inactivated single-stimulus trials (34 sessions in 6 mice), separately for different absolute stimulus contrast. Lines indicate average wheel velocity and shaded regions indicate 95% confidence interval. **(c)** Same analysis as in (b) but separated for high-contrast trials (54% contrast) with mixed-power optogenetic inactivation in left MOs (red), left MOp (orange) or with no inactivation (black). Gray arrow illustrates the decrease in peak velocity introduced by the laser inactivation. (**d**) Mean wheel velocity for open-loop (widefield imaging) and closed-loop (inactivation, laser off trials) experiments performed across separate experimental rigs. Colored lines indicate the average wheel velocity for each mouse, averaged over high-contrast (54% contrast) correct trials, and shaded regions indicate 95% confidence intervals. Gray shaded region indicates the time-window used for analyzing widefield and electrophysiological signals (Fig. 2). **(e)** Summary of the change in peak velocity across all mice, for Left (blue) and Right (orange) choices, for mixed-power inactivation in VIS, MOs, MOp and SSp in left or right hemispheres (text under plot indicates inactivated region). For each choice

and region inactivated, the percentage change in peak velocity is computed within each contrast condition and then averaged across conditions. Bars indicate the mean percentage change in peak velocity across mice and error bars indicate 95% confidence intervals. Significance across mice is determined by a t-test. Significance results are not corrected for multiple comparisons. *** p<0.001, ** p<0.01, * p<0.05. **(f)** Same analysis as in (e) but showing the percentage change in median reaction time for Left and Right choices. **(g)** Effect of high-power pulsed inactivation (15 mW, 25 ms) in VIS (blue) and MOs (green) on median reaction time (RT, pooling over 65 sessions in 6 mice) for correct trials. Same plotting convention as Fig. 3h. *** indicates time interval when median RT is significantly increased relative to control trials (*p*<0.0001; one-tailed Wilcoxon rank sum test). Horizontal dashed gray line: non-laser median RT. **(h)** Same as in (g) but for inactivation in MOp. The laser onset time on each trial was chosen randomly between -100 ms and +300 ms relative to stimulus onset on ~12% of trials (pooling data from 31 sessions in 6 mice). No significant effect of MOp inactivation was found for any time.

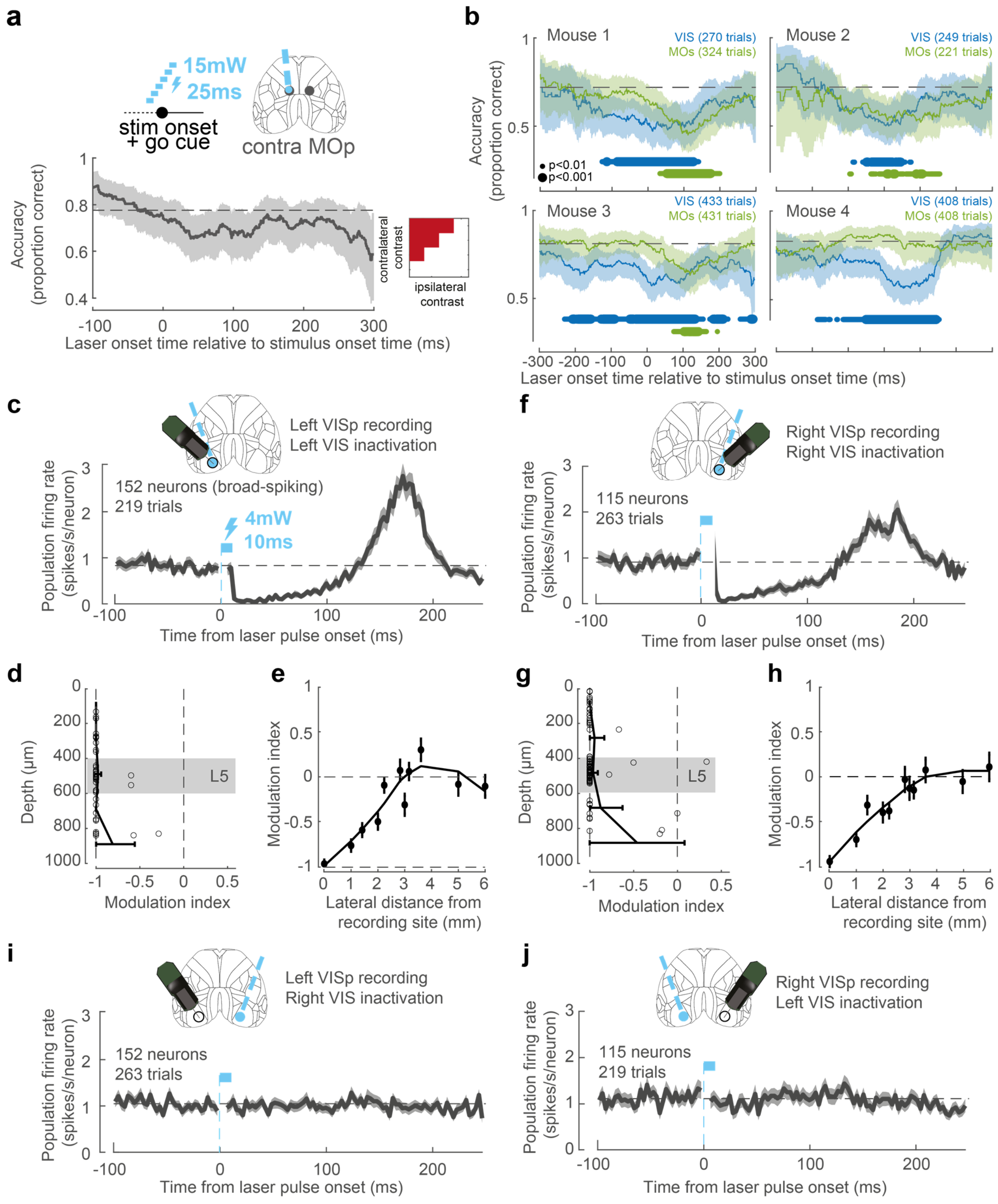


**Figure 3–figure supplement 3: Pulsed inactivation and electrophysiological recording. (a)** Pulse inactivation experiment in MOp. Graph shows performance (percentage correct, averaged over 100 ms boxcar window) as a function of laser onset time in MOp relative to stimulus onset. The laser onset time on each trial was chosen randomly between -100 ms and +300 ms relative to stimulus onset on ~12% of trials (25 ms pulse duration, 15 mW power, pooling data from 31 sessions in 6 mice). Dashed gray line: non-laser performance. No significant effect of MOp inactivation was found for any time. **(b)** Pulse inactivation experiment in VIS and MOs, shown for individual mice with at least 60 trials per inactivated region. Same plotting convention as Fig. 3h except using a 150 ms boxcar smoothing window. Colored dots indicate times when performance is significantly impaired relative to control trials (small dot: $p<0.01$, large dot: $p<0.001$; $\chi 2$ test). Number of inactivation trials in each region is shown inset. **(c)** Firing rate of 152 broad-spiking VISp neurons measured electrophysiologically in the left hemisphere, following a laser pulse (4 mW 10 ms) in the same region (grouped from multiple coordinates), in an awake mouse (Ai32 x PV-cre) not performing a task. The period during the light pulse is masked because recorded voltage deflections during this period may have corresponded to light artifacts. Error bars represent standard error across 219 trials. **(d)** Effect of inactivation by depth

from the cortical surface. Modulation index $(f - f_0)/(f + f_0)$, where $f_0$ is baseline firing (-100 to -25 ms) and $f$ is post-laser firing (+25 to +100 ms), is plotted for neurons in Left VISp spanning different cortical depths, following a laser pulse at the same location. Open circles represent single neurons, and error bars indicate the confidence interval of the modulation index for neurons binned by depth. Shaded region: approximate position of Layer 5 based on the density of neurons. **(e)** Spatial resolution of pulsed inactivation across cortical distance. Modulation index is plotted for Left VISp neurons recorded during pulsed laser illumination centered at different distances from the recording electrode. Dots and error bars indicate mean and 95% confidence interval of the modulation index across neurons. Black line is a LOESS smoothed curve. **(f-h)** Same as (c-e) but for 115 broad-spiking VISp neurons in the right hemisphere during pulsed inactivation in the right hemisphere. **(i,j)** Same as in (c,f) but shown for pulsed inactivation in the hemisphere opposite to the recorded hemisphere.

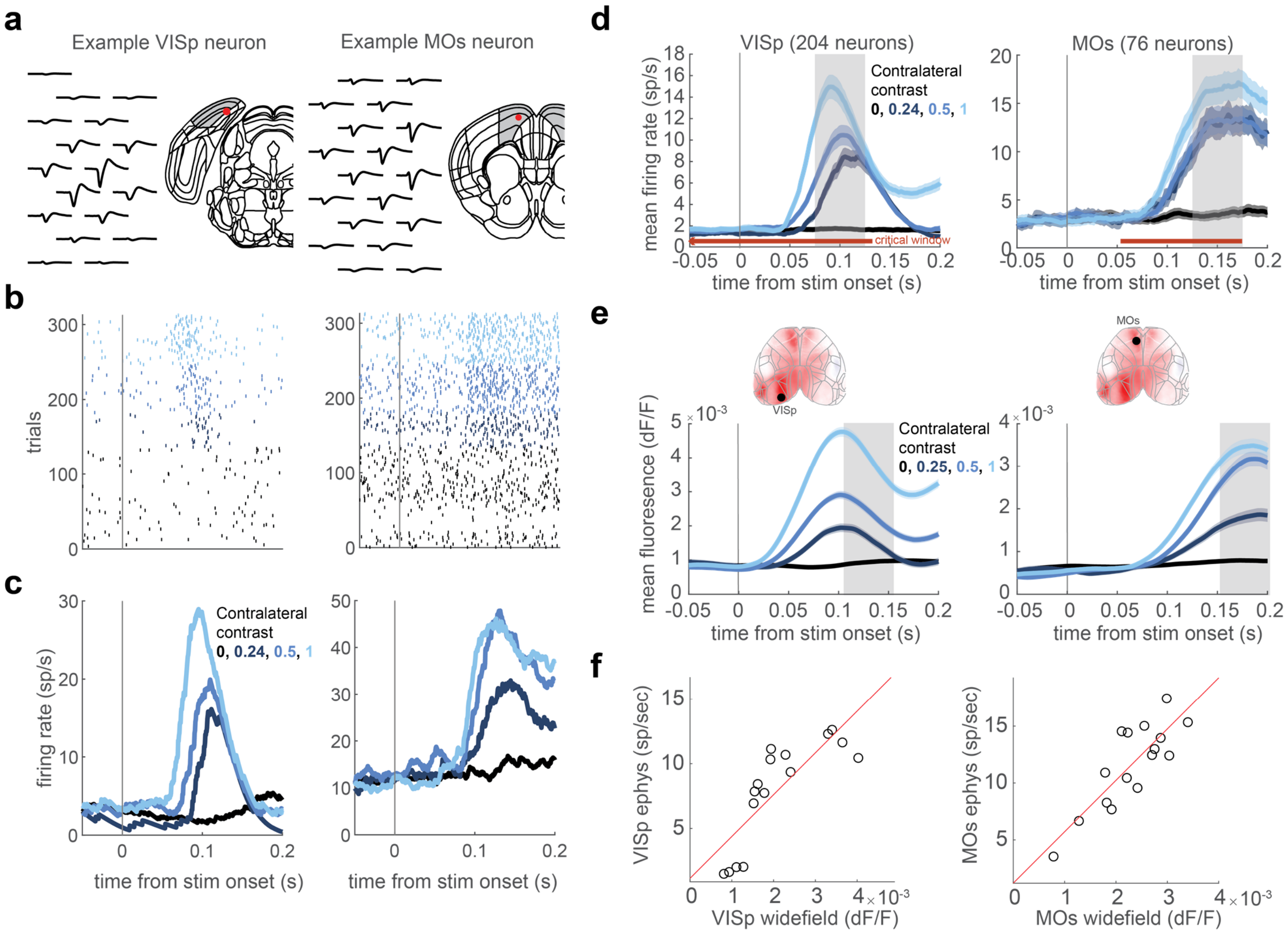


**Figure 4–figure supplement 1: Calibrating widefield activity to spike rate. (a)** Example neuron in Left VISp and Left MOs measured electrophysiologically. Mean waveforms are shown in black and the red dot marks the location of the neuron within an aligned Allen CCF atlas. **(b)** Raster plots showing spiking activity aligned to stimulus onset. Color reflects the contrast level presented to the contralateral hemifield (color code in panel c). **(c)** PSTHs for these example neurons. **(d)** Population PSTHs, averaged over 204 neurons in VISp and 76 neurons in MOs. Shaded color region marks the standard error across trials. The gray shaded regions mark the time window when the firing rate is averaged for subsequent analyses (VISp: 75-125 ms, MOs: 125-175 ms). Horizontal red line marks the time of the critical interval identified in the pulse inactivation experiment (Fig. 3h). **(e)** Same plotting convention as in (d) but showing trial-averaged widefield calcium fluorescence of Left VISp (Left) and Left MOs (right) ROIs in response to stimuli present on the contralateral side. Shaded regions mark the intervals used for averaging in subsequent analyses. This window is 30 ms after the window associated with electrophysiological data, to compensate for the slower kinetics of GCaMP6. Insets: ROI locations superimposed on mean fluorescence at 150 ms after stimulus onset. **(f)** Mean calcium fluorescence vs. population firing rates for Left VISp (left) and Left MOs (right). The 16 open circles correspond to the 16 contrast conditions. Red line corresponds to the fit of a simple linear model f(x) = b0 + b1*x. The linear model is used to transform widefield fluorescence data into estimates of population firing rate in the neurometric model (Fig. 4).

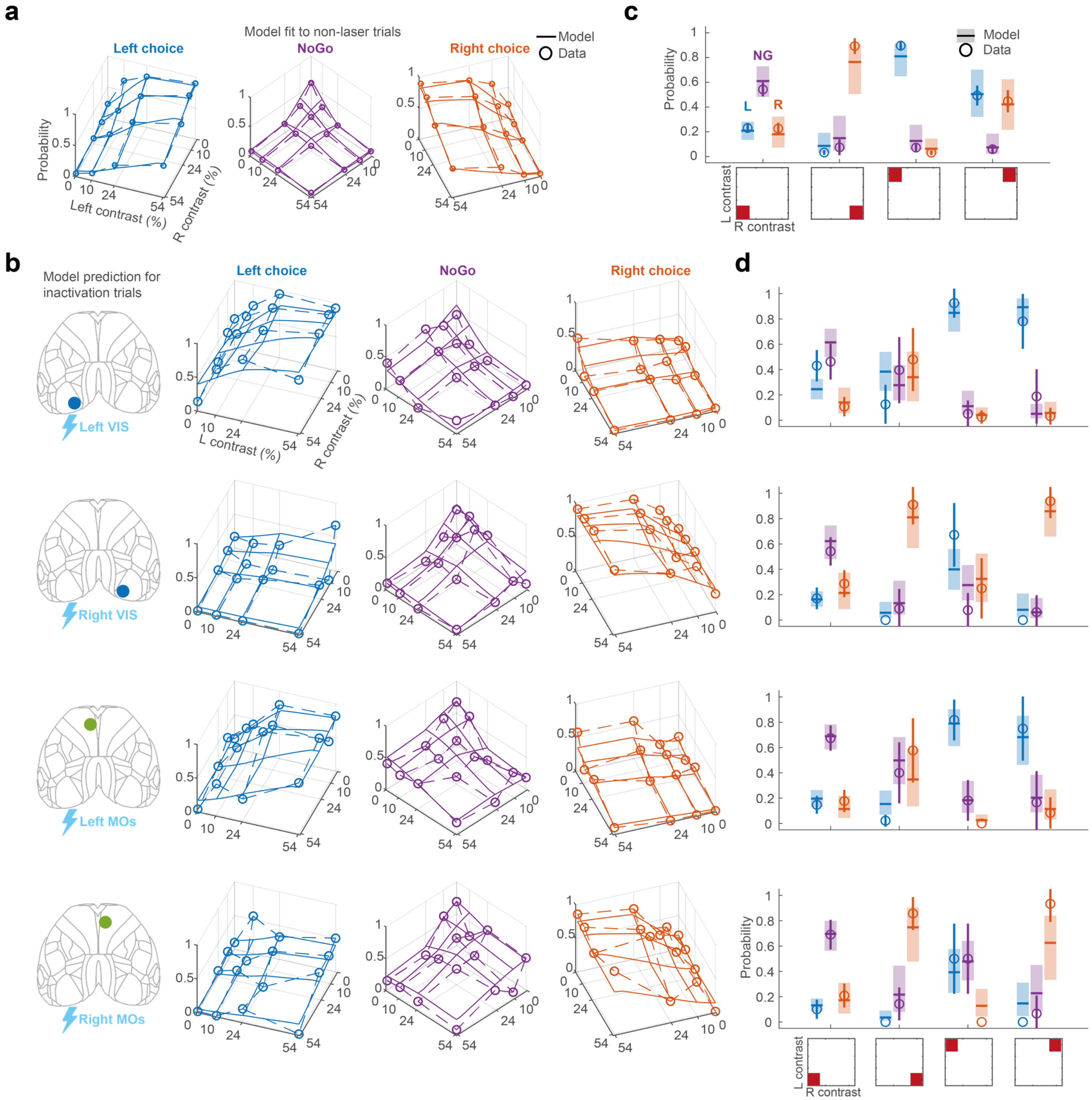


**Figure 4–figure supplement 2: Neurometric model fit and prediction. (a)** Fit of the neurometric model to non-laser trials from the mixed-power inactivation sessions, showing all contrast conditions on the left and right side. Same plotting scheme as in Fig. 1e-g. Empirical data are represented as colored dots for Left (blue), NoGo (purple) and Right (orange) choices, averaged across 34 sessions in 6 mice. Solid lines correspond to the posterior mean of the neurometric model fit. For each contrast condition, the probability of Left, NoGo and Right are computed using the fitted weights and average firing rate for each contrast condition (Figure 4-figure supplement 1) with interpolation between the contrast values tested. **(b)** Same plotting scheme as in (a) but showing the data and the prediction of the neurometric model when inactivating Left VIS, Right VIS, Left MOs and Right MOs for 1.5 s from stimulus onset, averaged over all laser powers. Model predictions are generated by setting the activity for each region to zero in the model. (**c**) Alternative visualization of the neurometric model fit to non-laser trials, focusing on trials with zero contrast, high contrast left or right stimuli, or high contrast bilateral stimuli (shown inset). Open circles indicate the empirical session-averaged proportion of Left (blue), NoGo (purple) and Right (orange) choices. Error bars are 95% confidence intervals. Colored lines and shaded region indicate the mean and 95% credible intervals of the posterior fit of the neurometric model. (**d**) Same as in (c) but showing the data and neurometric model prediction for each inactivated region corresponding to same row in panel (b).

[VIDEO FILE 1]

**Figure 2-video 1: Widefield fluorescence in VISp, MOs, MOp/SSp for different contralateral and ipsilateral contrast conditions.** Each panel shows the timecourse of fluorescence in one cortical region, for all 16 contrast conditions. Within each panel, the two black grids show the 40th and 60th percentile (across sessions) of widefield fluorescence (dF/F), averaged over all possible behavioral choices. Different frames of the video correspond to different times after stimulus onset. Time is indicated in the title of each panel.

[VIDEO FILE 2]

**Figure 2-video 2: Average widefield fluorescence in VISp, MOs, MOp/SSp for different contralateral and ipsilateral contrast conditions, and choices.** Similar to Figure 2-figure supplement 4, but dividing responses by behavioral choice. The three grids in each panel show mean cortical fluorescence as a function of contrast condition, separated for Contraversive (away from the ROI hemisphere; blue), NoGo (purple) and Ipsiversive (towards the ROI hemisphere; orange) choices. Conditions where there are fewer than 0.5 trials averaged across sessions (e.g. trials with high contrast on one side but the mouse moved the opposite direction) are excluded from plotting.

| | | | Session count for each Figure | | | | |
|---|---|---|---|---|---|---|---|
| **Name** | **Sex** | **Genotype** | **Fig. 2a-h** | **Fig. 2i-p *** | **Fig. 3a,d** | **Fig. 3g,h** | **Fig. 1e-h Fig. 4c,d** |
| Beadle | M | Ai32 x *Pvalb*-Cre | | | | 4 | 5 |
| Bovet | M | Ai32 x *Pvalb*-Cre | | | | 5 | 5 |
| Burnet | M | Ai32 x *Pvalb*-Cre | | | | 14 | 4 |
| Kornberg | M | Ai32 x *Pvalb*-Cre | | | | 14 | 8 |
| Medawar | M | Ai32 x *Pvalb*-Cre | | | | 14 | 4 |
| Ochoa | M | Ai32 x *Pvalb*-Cre | | | | 14 | 8 |
| Chomsky | F | Ai32 x *Pvalb*-Cre | | | 16 | | |
| Morgan | F | Ai32 x *Pvalb*-Cre | | | 8 | | |
| Murphy | M | Ai32 x *Pvalb*-Cre | | | 28 | | |
| Spemann | M | Ai32 x *Pvalb*-Cre | | | 26 | | |
| Whipple | F | Ai32 x *Pvalb*-Cre | | | 13 | | |
| Kendall | F | Ai95 x *Vglut1*-Cre | 2 | | | | |
| Moniz | M | Ai95 x *Vglut1*-Cre | 9 | 3 | | | |
| Muller | F | Ai95 x *Vglut1*-Cre | 1 | 3 | | | |
| Theiler | F | Ai95 x *Vglut1*-Cre | 1 | 1 | | | |
| Forssman | M | C57BL/6J | | 4 | | | |
| Richards | M | C57BL/6J | | 5 | | | |
| Chain | M | *Snap25*-GCaMP6s | 4 | | | | |
| Radnitz | F | *Snap25*-GCaMP6s | 6 | 5 | | | |
| Cori | F | tetO-GCaMP6s x *Camk2a*-tTA | 6 | 3 | | | |
| Hench | M | tetO-GCaMP6s x *Camk2a*-tTA | 7 | 4 | | | |
| Reichstein | M | tetO-GCaMP6s x *Camk2a*-tTA | 3 | | | | |
| Lederberg | F | *Vglut1*-IRES2-Cre-D | | 7 | | | |
| Tatum | F | *Vglut1*-IRES2-Cre-D | | 4 | | | |

**Supplementary Table 1: Mouse genotype and session count for each figure**

* Data previously published in Steinmetz, N.A., Zatka-Haas, P., Carandini, M., Harris, K.D., 2019. Distributed coding of choice, action and engagement across the mouse brain. Nature 1–8.
